# Host identity drives distinct prokaryotic and microeukaryotic microbiomes across four cold-water coral species in a near-pristine Mediterranean reef

**DOI:** 10.64898/2026.10.30.763042

**Authors:** A. Ardanaz-Ullate, M. Agazzi, C. Lo Iacono, J. del Campo

**Affiliations:** Institute of Evolutionary Biology (CSIC-UPF), Barcelona, Catalonia, Spain; Marine Sciences Institute (ICM-CSIC), Barcelona, Catalonia, Spain

**Keywords:** Cold water corals, DNA metabarcoding, microbiome, holobiont, 16S rRNA, 18S rRNA, Cabliers reefs, Alboran Sea

## Abstract

Cabliers is the widest Mediterranean cold-water coral (CWC) mound province exhibiting active growth. This area, a ridge-like CWC mound located in the Alboran Sea (SW Mediterranean) is characterized by thriving CWC reefs and accompanying mega-benthic communities (i.e.: gorgonians, black corals, sponges) found in near-pristine conditions and undergoing an almost null human pressure. We provide in this study the first characterization of the prokaryotic and microeukaryotic microbiomes of four key deep corals structuring the Cabliers province: *Madrepora oculata*, *Desmophyllum pertusum*, *Leiopathes glaberrima,* and *Acanthogorgia hirsuta*. We used DNA metabarcoding of the 16S and 18S rRNA gene V4 regions via Illumina MiSeq technology to elucidate symbiotic relationships that enable the holobiont to thrive. We found that host identity was the primary driver of microbial community structure, identifying specific signature microbiomes within coral species. The octocoral *A. hirsuta* showed reliance on microbes for organic matter processing such as the BD1-7 clade, while scleractinians *M. oculata* and *D. pertusum* displayed distinct symbionts, including *Endozoicomonas* (involved in the sulfur cycle) and the SUP05 clade, respectively. We suggest that coral microbiomes are vital for the continuous growth of Cabliers, with physiological needs and host identity shaping unique microbial communities within these deep coral ecosystems.

## INTRODUCTION

In the eastern Alboran Sea, the ridge-like Cabliers coral mound develops with a depth range of 290-700 m, along a N-S oriented 25 km long band (Corbera *et al*., 2019). It consists of several clusters of mounds, from few meters to tens of meters tall, hosting active and thriving cold-water coral reefs for around 6 kilometers in its northern sectors and soft coral communities and sponge fields laying on dead CWC frameworks in the central and southern sectors. Cabliers is the area where the highest aggregation of active scleractinian reefs is observed. Moreover, it remains barely altered by human activity, maintaining a near-pristine condition (Corbera *et al*., 2019; Lo Iacono *et al*., 2019). Unlike neighboring provinces such as the West Melilla mounds, Cabliers exhibits optimal, thriving biological activity and vertical reef frameworks while other spots are mainly composed of sediment and coral rubble (Lo Iacono *et al*., 2014). The biological and ecological uniqueness of Cabliers is primarily attributable to its near-pristine state of conservation. The interaction between inflowing fast and fresh Atlantic waters providing high surface productivity and the Mediterranean Levantine Intermediate Waters (LIW) with moderate currents, ensuring a regular flow of organic matter, provide the most suitable conditions for coral growth and a high associated diversity in Cabliers (Corbera *et al*., 2019, 2022).

The Cabliers reef is built by Cold Water Corals (CWC), a diverse group of cnidarians thriving between 200 and 2000 meters depth, without light and at temperatures close to 13°C or 14°C (Roberts *et al*., 2006; United Nations, 2017), survive by capturing suspended particles such as organic aggregates, phytoplankton remains or small crustaceans (Henry and Roberts, 2016). Often called “true ecosystem engineers” their skeletons serve as a refuge, nursery, and habitat for thousands of other species (Roberts and Cairns, 2014). Nevertheless, increasing anthropogenic activities, with specific regard to bottom trawling and deep longliners, and global changes such as temperature warming threaten these CWC ecosystems. The reef is mainly composed of four species of cold-water corals: *Madrepora oculata*, *Desmophyllum pertusum*, *Leiopathes glaberrima,* and *Acanthogorgia hirsuta*, spanning hard corals, black corals, and octocorals are the most dominant. Differences in their chemical composition, morphology (skeletons), and growth strategy differences (United Nations, 2017) likely drive distinct microbiome compositions among these species. *M. oculata* (*Scleractinia*) presents a rigid aragonite skeleton (Roberts *et al*., 2009) and represents almost 90% of the coral scleractinian species growing structuring the Cabliers reefs (Corbera et al., 2019), as for the other CWC provinces in the Mediterranean (Gori *et al*., 2023). *D. pertusum* is also a scleractinian coral and the most iconic coral reef builder in the Atlantic (Henry and Roberts, 2016), and recent genomic studies have shown that the survival of these populations is highly dependent on larval settlement preferences (Weinnig *et al*., 2024). *A. hirsuta* (*Octocorallia*) is a key gorgonian species. This coral forms dense, heavy coral gardens that are essential for associated fauna (Orejas and Jiménez, 2019), and it possesses a thick mucus layer and a flexible skeleton mainly composed of gorgonin (collagen-like protein), expected to host significantly different microbial life from scleractinians (stony corals) (Bo *et al*., 2014; Sartoretto and Zibrowius, 2018). As for other distinctive traits, *A. hirsuta* has been observed to accumulate metals such as copper and cadmium, making it an efficient bioindicator of environmental conditions (Raimundo *et al*., 2013). Finally, *L. glaberrima* (Antipatharia) differentiates itself from the other three corals by displaying a rigid yet flexible keratin-like (chitin and scleroprotein) skeleton and a longer lifespan, resulting in the ability to create dense forests (Massi *et al*., 2018; Lauria *et al*., 2021).

Recent discoveries connecting any organism’s health and its microbiome have also extended to corals. In consequence, the holobiont concept has grown in importance, defining this term as the whole symbiotic consortium formed by its associated microeukaryotic (fungi and protists) and prokaryotic (bacteria and archaea) communities (van Oppen and Blackall, 2019). These are not merely inhabitants, as it has recently been shown that the coral microbiome has a relevant role in sickness and health (Voolstra *et al*., 2024). Therefore, coral holobiont can be considered a further important factor contributing to the optimal state of conservation and active growth of CWC reefs (Hernandez-Agreda *et al*., 2017). And not all its members are equally prevalent. We define the concept of core microbiome as the stable and most persistent fraction of the microbiome, sustaining relevant symbioses (Hernandez-Agreda *et al*., 2017). This includes functional microorganisms such as bacteria, viruses, archaea, fungi, and protists (van Oppen and Blackall, 2019). Thus, it is crucial to elucidate the mechanisms underlying the ecological success of this population. Cnidarians are holobiont organisms whose physiological stability and survival depend on their associated microbiome. The main focus of this project is to characterize this symbiotic network, which has never before been described from this unique enclave. To capture this microbial complexity, we employed DNA metabarcoding. A high-throughput sequencing method that identifies multiple taxa from a single biological sample; thus, it is the most precise technique for unveiling the Cabliers microbiome (del Campo *et al*., 2026).

Taking together, we assessed the taxonomic and functional diversity expected within the microbial consortium, and the methodological tools capable of capturing it. Metabarcoding becomes the tool enabling us to characterize the full symbiotic community shaping the health and resilience of each species. By comparing the microbial fingerprints of *A. hirsuta*, *M. oculata*, *D. pertusum*, and *L. glaberrima*, this approach will provide a first baseline description of the Cabliers holobiont, a step toward a better understanding of reef dynamics.

## EXPERIMENTAL PROCEDURES

### Sample collection

The Cabliers CWC mound province, located in the eastern Alboran Sea (SW Mediterranean) between 35°39′–35°48′ N and 2°14′–2°18′ W, consists of ridge-like mounds up to 70 m high clustered along a N–S direction for approximately 25 km. Living reefs occur along the crests of the mounds in the northern sector, whereas the rest of the province is occupied by well-preserved, long-lived structuring communities, mainly soft corals (gorgonians and antipatharians) and sponge fields.

Coral samples from *A. hirsuta*, *M. oculata*, *D. pertusum* and *L. glaberrima* (chunks of 1cm average) were collected in Cabliers during the 2^nd^ to the 25^th^ of April 2024 (PROTEUS cruise of the PROTEUS Spanish National Project, R/V *Sarmiento de Gamboa*) using the work-class remotely operated vehicle (ROV) Luso (EMEPC, Portugal) in six exploratory dives (dives 2, 5, 9, 11, 14 and 17) within a depth range of 290–400 m. The two six-function hydraulic manipulators of the ROV allowed precise sampling of each specimen. Samples were stored in DNA Shield (Zymo Research) at ambient temperature on board and at 4°C from their arrival at the laboratory until November 2025. Sampling covered the whole 25 kilometers long CWC system, not only the northern sector where living reefs occur. Quadruplicate samples were collected to provide backups and support a triplicate system approach. Resulting in 39 DNA samples (*A. hirsuta*; n = 9, *M. oculata*; n = 9, *D. pertusum*; n = 12, and *L. glaberrima*; n = 9).

### DNA extraction and quantification

We extracted triplicates of each sample type (species per location) using the ZymoBIOMICS DNA/RNA Miniprep Kit following a specific workflow due to the nature of our samples. All coral biopsies that were preserved in DNA Shield were grounded with mortar an pestle in ice, then they underwent extended mechanical lysis by high-speed bead beating for a total of 20 minutes to ensure maximum yield from the calcium carbonate skeleton and coral tissues. Then, the supernatant was treated with the specific kit lysis buffer (2:1), and genomic DNA (gDNA) was purified using a spin-away filter, while the flow-through contained RNA. We precipitated the flow-through with pure ethanol, treated it with DNase I, and purified it by column (Zymo spin). DNA and RNA were purified using Zymo spin HRC (inhibitor removal) filters to remove PCR inhibitors, and finally, they were eluted in nuclease-free water to a final volume of 60 µl. After extraction, DNA yield was quantified using the Qubit High Sensitivity DNA Kit and met the sequencing requirements for a 20 µl sample with a minimum concentration of 50 ng/µl. After extractions, DNA was stored at 4°C prior to PCRs and then at −20°C prior to shipping with cold bags to the sequencing facility at Dalhousie University – IMR (Integrated Microbiome Resource) in Canada.

### 16S and 18S rRNA genes Amplification and Sequencing

To characterize the microbial communities associated with coral samples, we amplified the V4 region of the 16S and 18S genes to identify prokaryotes and eukaryotes, respectively. Moreover, to sequence our 39 samples, we employed Illumina MiSeq technology at the Dalhousie University IMR (Integrated Microbiome Resource) in Canada. We proceeded with DNA amplification using GoTaq DNA Polymerase, with antimetazoan primers forward (574F) and reverse (UnNonMet DB) (del Campo *et al*., 2019) in order to prune out all the metazoans from our DNA and to overexpress the DNA of all other eukaryotes. All reactions were prepared in a final volume of 25 µl, following the manufacturer’s instructions and under the following conditions: 98 °C for 30 s; 34 cycles of 98 °C for 10 s, 51.1 °C for 30 s and 72 °C for 60 s; and a final extension at 72 °C for 5 min. Then, we proceeded with a second PCR targeting the 18S only; this second PCR was performed by the sequencing facility. Library preparation was performed by the sequencing facility using the standard Illumina protocol. For the prokaryotic community, we submitted genomic DNA to IMR. We sent a total volume of 20 µl per sample with a minimum concentration threshold of >50 ng/µl. DNA was amplified using the following universal primers: Forward (515F) and reverse (806R) (Apprill *et al*., 2015; Parada *et al*., 2016). The sequencing facility performed the PCR amplifications, and sequencing was conducted using a Paired-End 250 bp (PE250) configuration. For assessing the microeukaryotic community, 20 µl of PCR products per sample were submitted instead of genomic DNA. Prior to submission to the IMR, an antimetazoan PCR was performed with the following pair of primers in order to exclude metazoan templates: forward (574F) and reverse (UnNonMet DB). The products of this first PCR were used by the sequencing facility as templates for the second PCR, targeting the V4 region of the 18S rRNA gene with the following primers: forward (572F) and reverse (E1009R) (Comeau *et al*., 2011). IMR performed the PCR amplifications, and sequencing was conducted using a Paired-End 300 bp (PE300) configuration. However, host reduction was only partially effective: 99.6% of the 18S rRNA gene reads were assigned to Metazoa (98.9% to Cnidaria), so the number of reads available for the microeukaryotic community after filtering was limited (66 to 1362 reads per sample; Appendix A. Table S3).

### Microbiome analysis

An average of 42,116 16S reads per sample (median: 41,310) and 126,318 18S reads per sample (median: 127,466) were recovered. Primers, adaptors, and low-quality bases were removed using Cutadapt v4.4 (Martin, 2011) and then, paired-end reads were processed in R using the DADA2 package (v1.38.0) (Callahan *et al*., 2016a). According to their quality profiles, on the one hand, 16S forward reads were truncated at 250 bp and reverse reads at 200 bp and filtered using the parameters maxN=0, maxEE=c(2,2) and trimLeft=10 because, based on empirical observations, it has been seen that on Illumina datasets these positions are more likely to retain pathological errors (Callahan *et al*., 2016b). On the other hand, 18S forward reads were truncated at 280 bp and reverse reads at 210 bp, and filtered using the parameters maxN=0, maxEE=c(2,5), and trimLeft=10. Then, for both sample types, paired-end reads were denoised and merged, chimeras were removed, and reads were grouped into amplicon sequence variants (ASVs) **(Appendix A. Tables S1-S2)**. Next, prokaryotic ASV taxonomy was assigned using the SILVA v138.1 database as a reference (Quast *et al*., 2013), and for eukaryotes, the PR2 18S rRNA database v4.14.0 (Guillou *et al*., 2013) was used. The resulting ASVs tables, the taxonomic assignments, and the metadata for both sample types (16S and 18S) were integrated into a single unit of analysis, the phyloseq object, using the phyloseq package (McMurdie and Holmes, 2013), obtaining a total of 6792 ASVs for the 16S and 2103 ASVs for the 18S with an overall of 39 samples for both of them **(Appendix A. Tables S1-S2) (Appendix A.)**. *Chloroplast*, *Mitochondria*, *Metazoa, Embryophyceae* and contaminants were removed. In addition, for the 16S dataset, only ASVs that represented the top 99% of reads in at least two samples were retained, and samples with fewer than 250 reads were discarded. In contrast, for the 18S dataset, only the 99% relative abundance filter was applied, as we were trying to capture the most microeukaryotic diversity due to how predominant prokaryotes in non-photosynthetic coral microbiomes are (Mayer *et al*., 2025). We obtained a final filtered phyloseq object with 4074 ASVs for 16S and 391 ASVs across 39 samples for both sample types. Retrieving the following reads per sample **(Appendix A. Table S3)**. For the microbiome analysis, alpha diversity was assessed using the microbiome R package (https://microbiome.github.io/) and calculated with the Shannon index, and differences between species were tested with pairwise Wilcoxon rank-sum tests. Community beta diversity was examined using Aitchison distances, and ANOSIM with 999 permutations (Andersen *et al*., 2018). By utilizing center log ratio (CLR) transformations (Gloor *et al*., 2017), we proceeded with Principal Component Analysis (PCA). Then, for compositional analysis, UpSet plots, bubble plots, and heat maps were performed. Core microbiomes were developed with plot core by plotting, for the ten most abundant ASVs of each species, their prevalence across ten logarithmically spaced detection thresholds from 0.01% to 20% relative abundance. Furthermore, figures were constructed using ggplot2 and tidyverse packages (Wickham *et al*., 2019), and statistical analyses were done using vegan (Oksanen *et al*., 2025). Finally, the analysis of Compositions of Microbiomes with Bias Correction (ANCOM-BC2) was used to identify significantly enriched taxa among the four coral species (Lin and Peddada, 2020, 2024).

## RESULTS

### Alpha-diversity

Prokaryotic microbiome alpha-diversity analysis showed significant differences among some of the four coral communities studied in this project. Specifically, *A. hirsuta* showed the most stable and least diverse microbial structure, as indicated by the small Shannon Diversity Index (SDI), with significantly lower prokaryotic SDI than *D. pertusum* (p=0.018) and *L. glaberrima* (p=8.2·10^-5^) **(Figure 1A,I)**. *L. glaberrima* maintained consistently high, stable diversity, whereas *M. oculata* showed high inter-variability and the lowest diversity, but none of their comparisons were significant. Regarding the eukaryome alpha-diversity analysis, we observed an inverse pattern. Whereas *A. hirsuta* showed one of the lowest SDI for the prokaryotic community, in this case, this species presented the highest SDI, showing the most diverse microeukaryotic diversity with a median of 4.49, finding significant microbial community differences between all the other three corals (*D. pertusum* (p=6.8·10^-6^), *L. glaberrima* (p=4.1·10^-5^) and *M. oculata* (p=0.00016)). Moreover, we find significant differences between *D. pertusum* and *L. glaberrima* (p=0.041), indicating that *D. pertusum* has a significantly higher microeukaryotic diversity. *M. oculata* was more consistent in its microeukaryotic diversity than in its prokaryotic one, finding a significantly higher diversity of microeukaryotic (median=2.82) than in the case of *L. glaberrima* (p=0.05). Finally, there were no significant microeukaryotic SDI differences between *D. pertusum* and *M. oculata* (p=0.7) **(Figure 1A,II)**.

**Figure 1.**
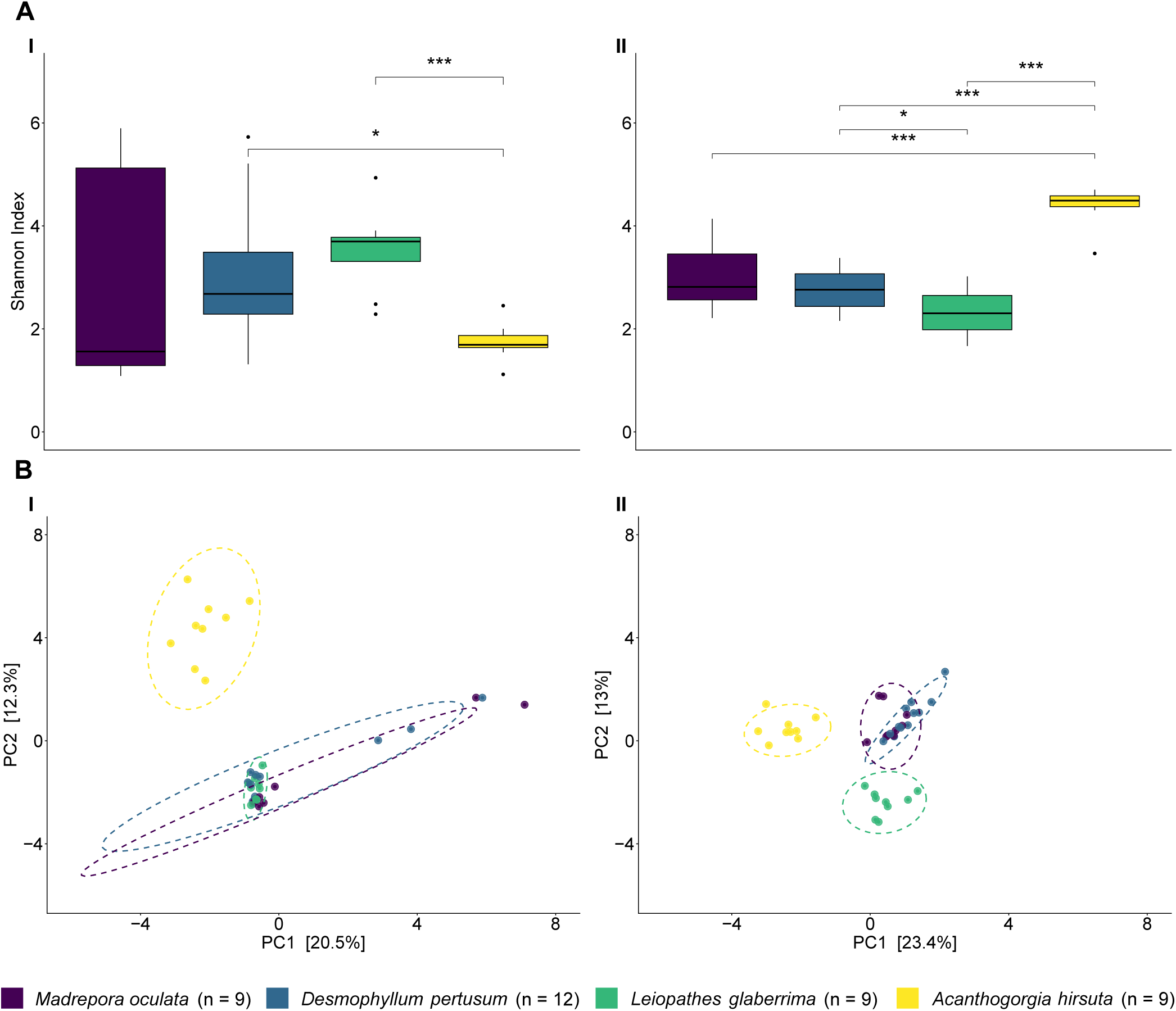
Alpha and beta diversity of prokaryotic and microeukaryotic communities across the four studied coral species. (A) Shannon diversity index for (I) prokaryotic and (II) microeukaryotic communities; asterisks indicate significant pairwise differences (padj ≤ 0.05; * padj ≤ 0.05, *** padj ≤ 0.001). (B) Principal Component Analysis (PCA) based on Aitchison distances for (I) prokaryotic and (II) microeukaryotic communities, with ellipses representing the 95% confidence interval per species. Beta diversity differed significantly among species for both the prokaryotic (ANOSIM, 999 permutations, R = 0.202, p = 0.001) and microeukaryotic (ANOSIM, 999 permutations, R = 0.486, p = 0.001) communities. Colors identify coral species, and n indicates the number of samples per species.

### Beta-diversity

Analysis of Similarities (ANOSIM) was used as the statistical test to identify differences in the microbial community among our target species. For both the prokaryotic (ANOSIM; 999 permutations, R = 0.202, p = 0.001) and the microeukaryotic (ANOSIM; 999 permutations, R = 0.486, p = 0.001) **(Appendix B. Figure S3)** communities, we found that species identity was a major driver of dissimilarity. In the PCA, the prokaryotic communities of *D. pertusum*, *L. glaberrima*, and *M. oculata* clustered together, while *A. hirsuta* showed a significant lower similarity to them and separated from the group. Moreover, *L. glaberrima* showed the lowest variability in terms of community structure **(Figure 1B,I)**. For the eukaryome beta diversity analysis, we found more separated communities Not only *A. hirsuta* was significantly different than the other three corals, but also *L. glaberrima* presented a significantly different microeukaryotic community compared to the other species, in spite of the fact that it showed greater variability than for the prokaryotic community. Lastly, *M. oculata* and *D. pertusum* presented a similar microeukaryotic community structure **(Figure 1B,II)**. In contrast to the identity of the corals, the variable “dive” (different dives were samples were collected) did not appear to be a major driver of dissimilarity for the prokaryotic beta diversity analysis (ANOSIM; 999 permutations, R = 0.1, p = 0.096), but not for the microeukaryotic community (ANOSIM; 999 permutations, R = −0.012, p = 0.548) **(Appendix B. Figures S3 and S4)**.

### Prokaryotic and microeukaryotic compositional analysis

Prokaryotic communities were dominated by Proteobacteria in all four corals, mostly within the order Pseudomonadales **(Figure 2A)**. Nitrosopumilales was the only abundant archaeal order, and it was present in every species. *M. oculata* and *D. pertusum* showed a diverse composition; however, there is also a clear dominance of Proteobacteria. For the other two corals, *A. hirsuta* showed the least diverse community, and it was also the most disctint: finding Entomoplasmatales from the Firmicutes phylum, not present in the other corals **(Figure 2A)**. *L. glaberrima,* unlike the other species, did not show Verrucomicrobiota, and it was also dominated by Proteobacteria_IS **(Figure 2A)**. For the microeukaryotic community, at the subdivision level, the composition was found to be mainly composed of Dinoflagellata, Radiolaria, and Ciliophora **(Figure 2B)**. Within the Dinoflagellata, the microbiome across all species was dominated by the orders Ichthyodinida, Syndiniales, and Gymnodiniales. Abundant microeukaryotic orders of the Radiolaria included Spumellaria and Acantharea, highlighting Acantharea_F only for *A. hirsuta* and *M. oculata,* or Acantharea_B for *L. glaberrima,* and RAD-B_IS just for *M. oculata*. Within Ciliophora, Apostomatia dominated *L. glaberrima* microbiome, whereas Choreotrichida and Suctoria dominated the other three corals **(Figure 2B)**.

**Figure 2.**
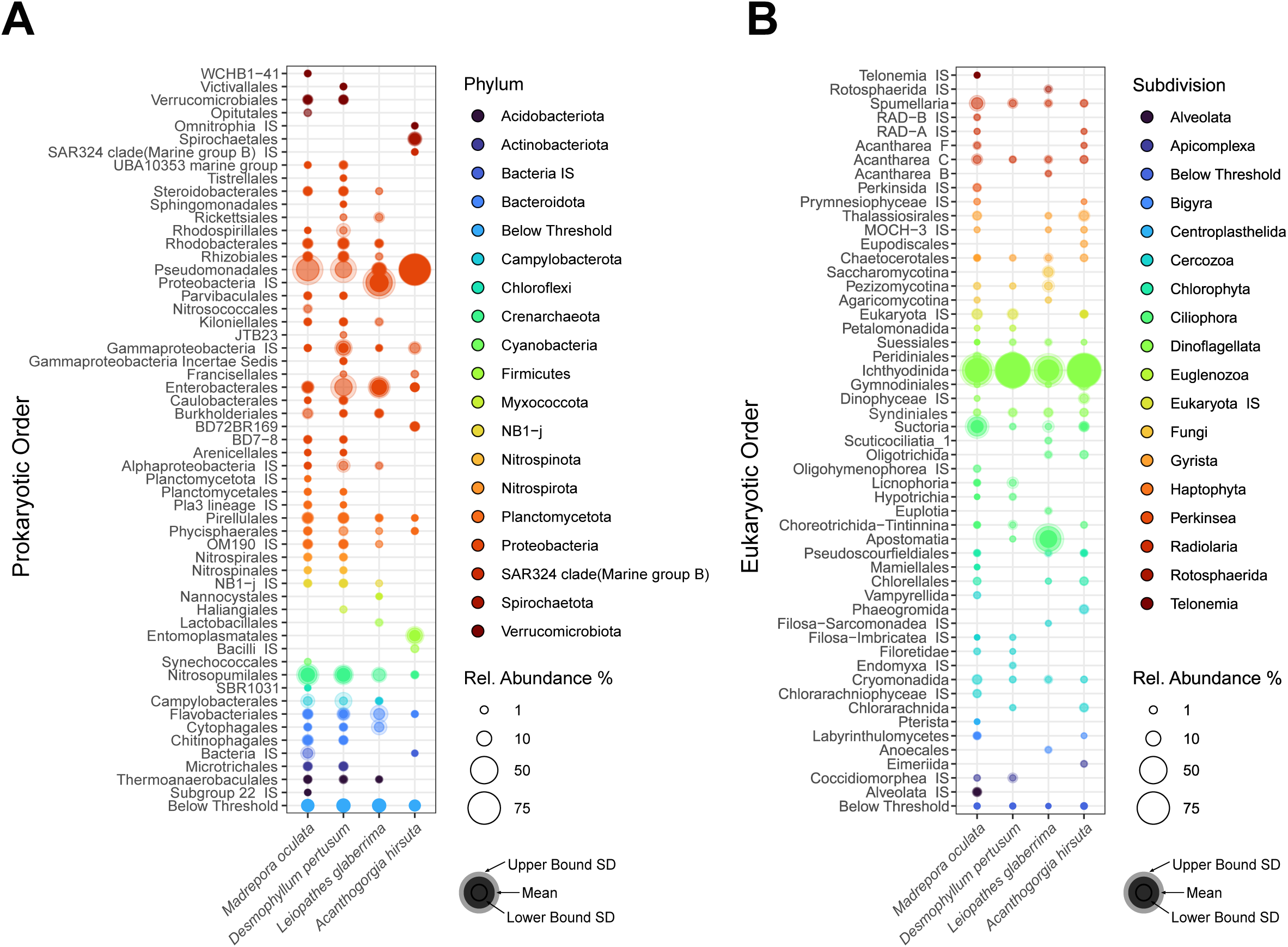
Relative abundance of the most abundant (A) prokaryotic and (B) microeukaryotic orders in the microbiome of the studied corals. Circle size is proportional to relative abundance; color indicates phylum (A) or subdivision (B). Only taxa with a median relative abundance >0.25% (prokaryotes) or >0.05% (microeukaryotes) are shown; remaining taxa are grouped as “Below Threshold.” The outer, semi-transparent bubble represents the upper bound of the standard deviation; the solid inner circle denotes the lower bound. Syndiniales_Group-I corresponds to Ichthyodinida. IS: *incertae sedis*.

To resolve community structure at a finer level, we generated heat maps of the most abundant amplicon sequence variants (ASVs). For the prokaryotic community, three of the four corals were each dominated by a single, species specific ASV: *Endozoicomonas* ASV_5 in *M. oculata* (37.8% relative abundance), SUP05 cluster ASV_14 in *D. pertusum* (32.2%), and BD1-7 clade ASV_2 in *A. hirsuta* (65.2%). All three belonging to the Proteobacteria phylum. *A. hirsuta* showed also a 10.6% read abundance from *Hepatoplasma* (*Candidatus Hepatoplasma* and *Candidatus Oceanoplasma*). *L. glaberrima* showed a different ASVs structure within Proteobacteria. Its community was dominated by Proteobacteria_IS ASV_7 (17.7% non-identified taxa), *Blattabacteriaceae*_IS ASV_71 (8.2%), *Candidatus Amoebophilus* ASV_84 (8.0%), and *Alteromonas* ASV_51 (6.9%). Among the most abundant ASVs, none was shared between all coral species. Nevertheless, *Nitrosopumilus* was the only genus consistently present in every species (ASVs 8, 10 and 11) **(Figure 3A)**.

**Figure 3.**
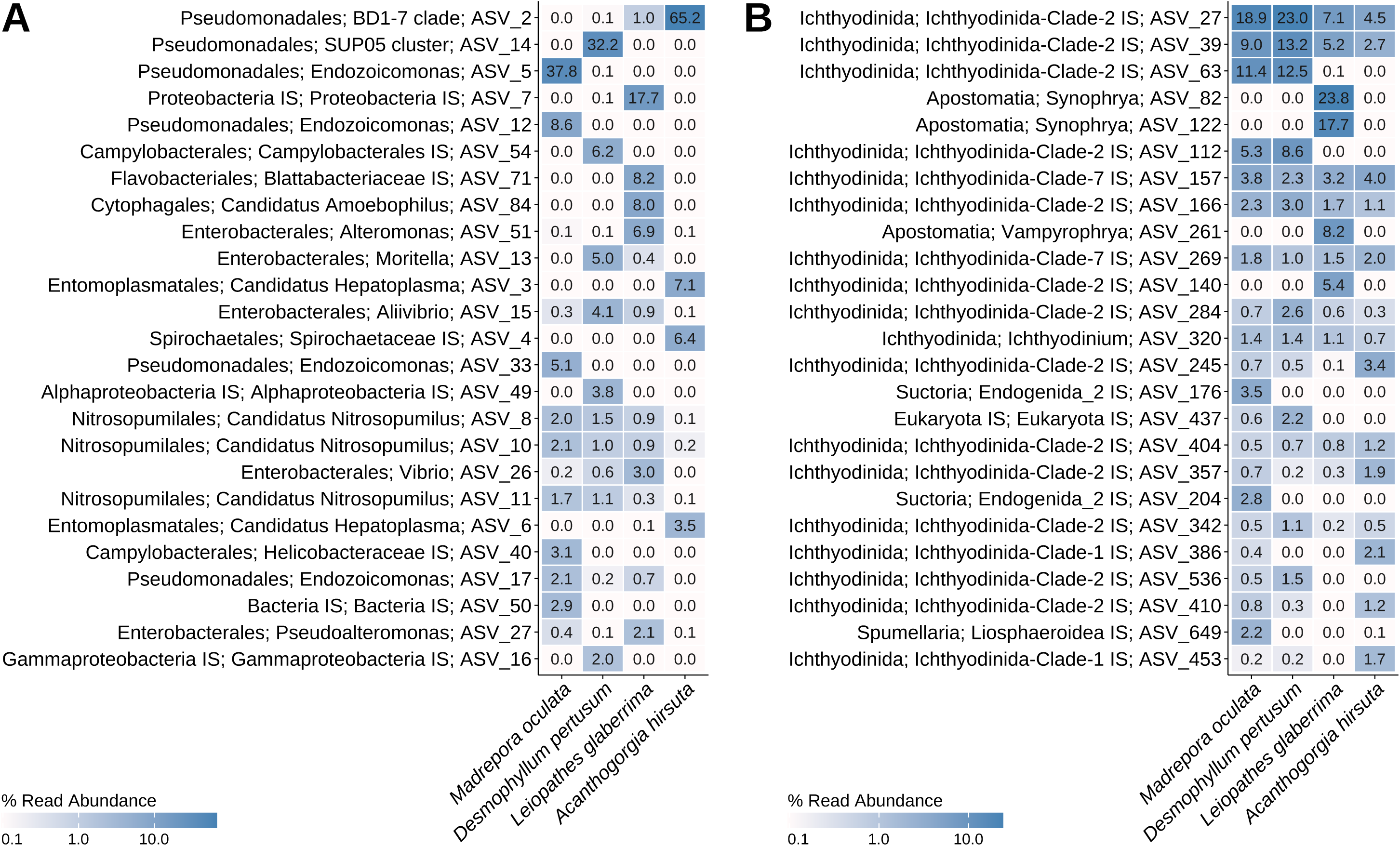
Mean relative read abundance (%) of the 25 most abundant (A) prokaryotic and (B) microeukaryotic ASVs in each coral species. ASVs are labeled as order; genus; ASV, and the value in each cell indicates the mean percentage of reads of that ASV in the species. The color gradient (white to blue, logarithmic scale) is specific to each panel. IS: *incertae sedis*.

For the microeukaryotic community, the results were similar across species; the eukaryome is dominated by the Dinoflagellata subdivision, specifically by Ichthyodinida-Clade-2 and Clade-7 species. However, *L. glaberrima* again stood out. It harboured the least diverse microbial community, dominated by a 41.5% read abundance of *Synophrya*, followed by *Vampyrophrya* (8.2% of their total read abundance), both from the Ciliophora subdivision, and neither was detected in the other three coral species **(Figure 3B)**. *A. hirsuta*, *M. oculata* and *D. pertusum* were dominated by Ichthyodinida-Clade-2 and Clade-7 species, read abundances were more similar for *M. oculata* and *D. pertusum*. In fact, ASVs 27, 39, and 63 read abundances (Ichthyodinida-Clade-2) from *M. oculata* and *D. pertusum* are higher than those presented in *A. hirsuta* and *L. glaberrima*.

After identifying the dominant ASVs for each species and community, we next determined how many were exclusive and how many were shared between species. For the prokaryotic community, 1087 ASVs were shared among the four species. *A. hirsuta, D. pertusum,* and *M. oculata* shared 108 ASVs, and only between *D. pertusum* and *M. oculata* were 688 ASVs shared. Moreover, *M. oculata* showed the highest number of exclusive ASVs, while *L. glaberrima* just had 28 exclusive ASVs **(Figure 4A)**.

**Figure 4.**
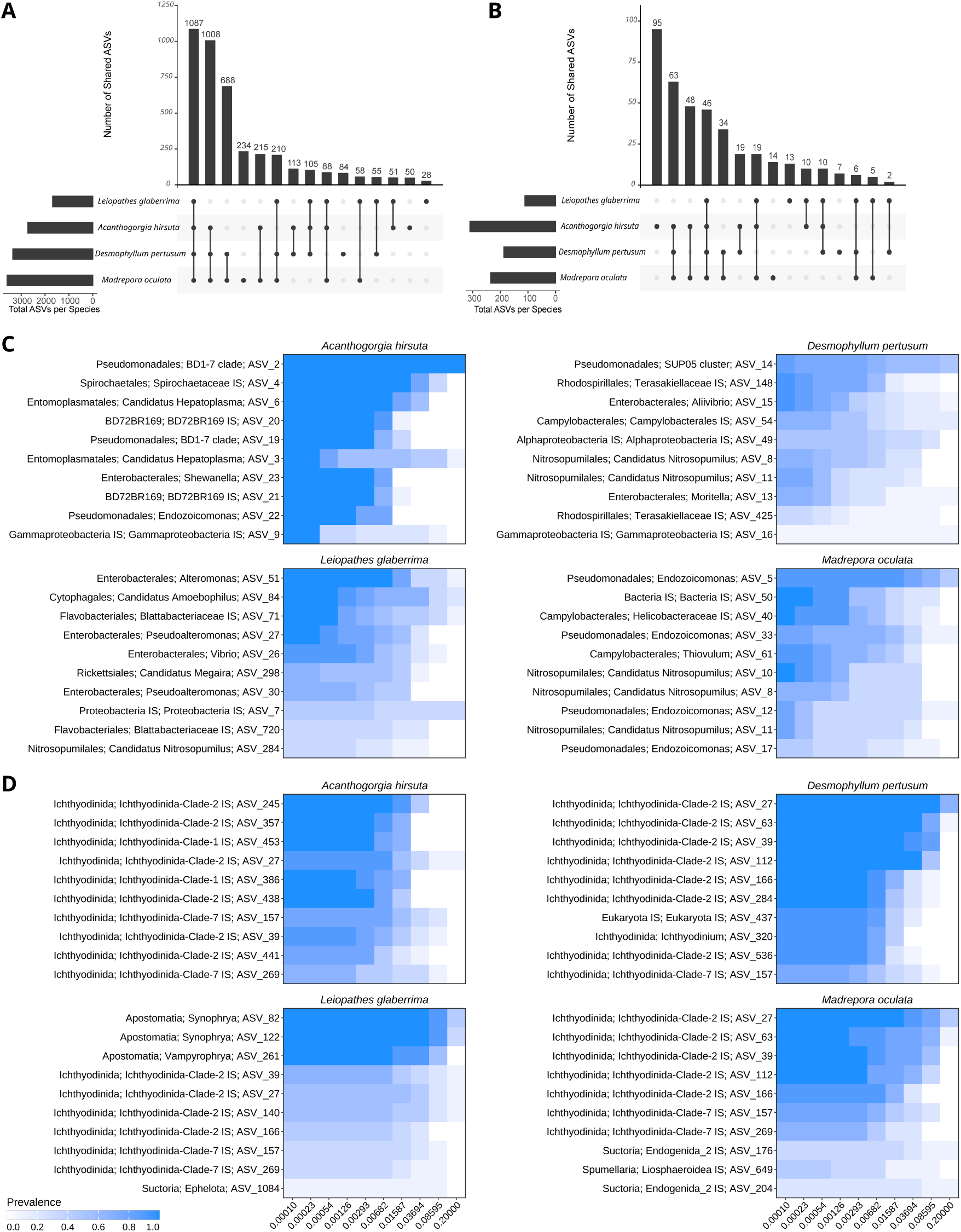
Shared and core microbiomes of the four coral species. (A, B) UpSet plots showing the number of ASVs shared among species or exclusive to one species for the (A) prokaryotic and (B) microeukaryotic communities; horizontal bars indicate the total number of ASVs per species and vertical bars the number of ASVs in each intersection, defined by the connected dots. (C, D) Core microbiome of each species for the (C) prokaryotic and (D) microeukaryotic communities, shown as prevalence heatmaps across ten logarithmically spaced detection thresholds from 0.0001 (0.01%) to 0.2 (20%) relative abundance. Color indicates prevalence, that is, the proportion of samples of each species in which the ASV exceeds the detection threshold (white, 0; blue, 1). Up to ten of the 25 most abundant ASVs of each species are shown, labeled as order; genus; ASV. IS: *incertae sedis*.

Among the microeukaryotic community, *A. hirsuta* had the highest number of exclusive ASVs, whereas *D. pertusum* had only 7 ASVs. As in the prokaryotic analysis, A. *hirsuta, D. pertusum,* and *M. oculata* shared the highest number of ASVs, with a substantial overlap between *D. pertusum* and *M. oculata* **(Figure 4B)**.

#### Core microbiome

In addition to UpSet plots, we identified the exclusive ASVs for each species and sample type (16S or 18S), visualized them via heatmaps, and found that ASVs with significant percentage read abundance were present in the species core microbiomes.

Within the prokaryotic core microbiome, *A. hirsuta* showed absolute prevalence and a high relative abundance of the BD1-7 clade, appearing in 100% of samples with at least 20% relative abundance. This coral showed the most different core microbiome. *M. oculata* and *D. pertusum* presented a similar core microbiome dominated by the phyla Proteobacteria, Crenarchaeota, and Campylobacterota, highlighting *Endozoicomonas* for *M. oculata* and the SUP05 cluster for *D. pertusum*. *L. glaberrima* showed the most diverse core microbiome, but without ASVs above 2.52% of read abundance. Moreover, we found *Nitrosopumilus*, *Aliivibrio*, and *Pseudoalteromonas* in every species’ core microbiome. Besides, *Endozoicomonas* was also found in the *A. hirsuta* core microbiome **(Figure 4C)**.

With respect to the core microbiome analysis of the microeukaryotic community, Ichthyodinida-Clade-2 dominated every coral core microbiome. However, for *L. glaberrima*, as shown in the upset plot, we also identified 7 exclusive ASVs, with a core microbiome dominated by *Synophrya* and *Vampyrophrya*. Moreover, we found *Ichthyodinium* in all coral core microbiomes, except in *A. hirsuta* **(Figure 4D)**.

### Differential abundance

#### ANCOM-BC2 analysis

At this point, we knew the main ASVs and their distribution across coral species; we wanted to know whether taxon abundance differed significantly among species and whether these differences were associated with dominant ASVs. To do so, we ran ANCOM2 with one-versus-all comparisons, visualized results with volcano plots **(Figure 5)** and bar plots **(Appendix B, Figure S4)** and labeled the top five significant taxa at the ASV level.

**Figure 5.**
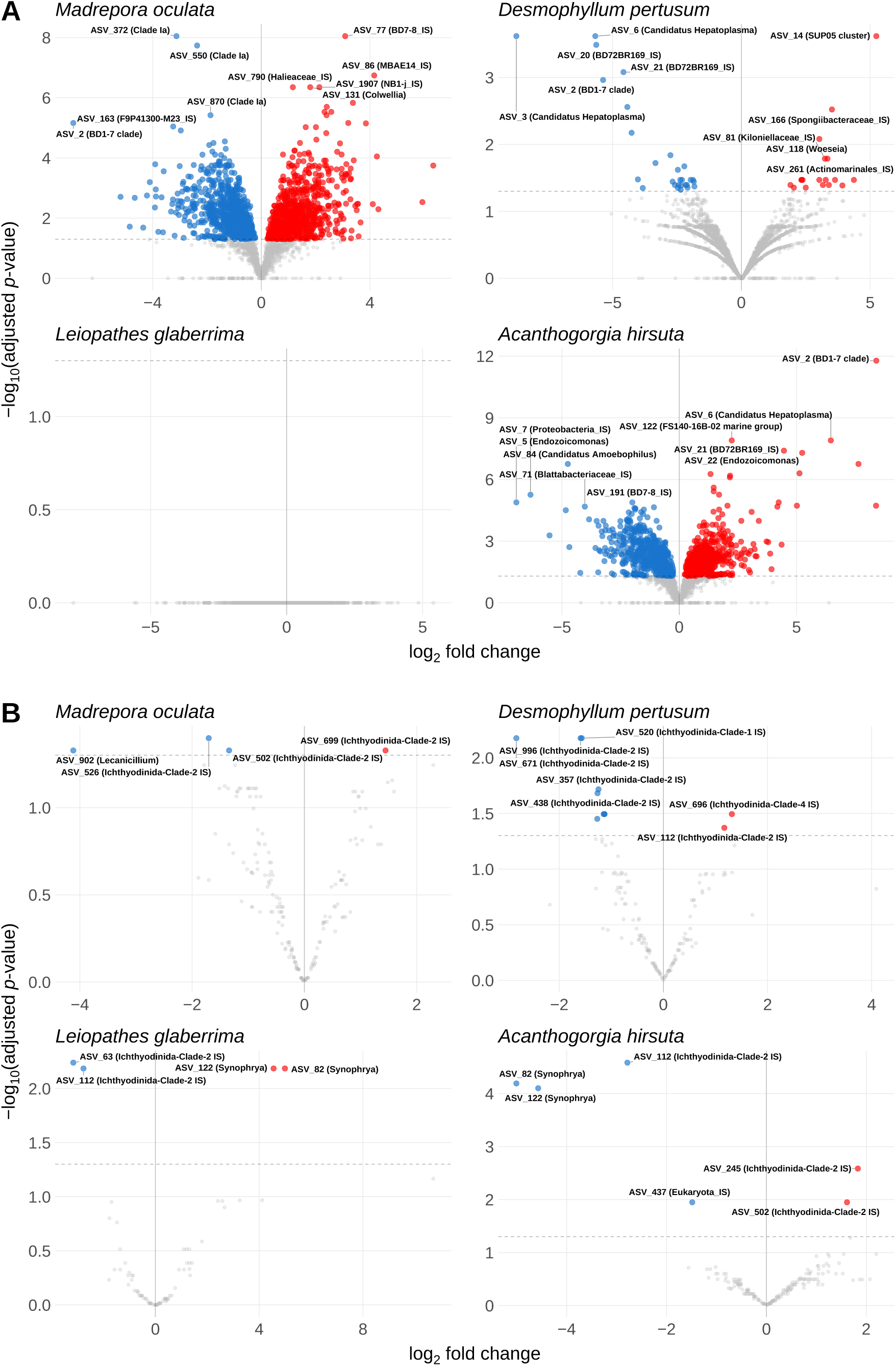
Differentially abundant ASVs in each coral species compared with the other three species (one-versus-all ANCOM-BC2) for the (A) prokaryotic and (B) microeukaryotic communities. Each point is an ASV; the x-axis shows the log_2_ fold change and the y-axis the −log_10_ of the FDR-adjusted p-value. Significantly enriched ASVs (q < 0.05, positive fold change) are shown in red, significantly depleted ASVs (q < 0.05, negative fold change) in blue and non-significant ASVs in grey; the dashed line marks q = 0.05. The five most significant enriched and depleted ASVs are labeled with their genus or lowest assigned taxonomic rank. No prokaryotic ASV was significantly enriched or depleted in *L. glaberrima*. IS: *incertae sedis*.

Regarding the prokaryotic community, *A. hirsuta,* was differentially enriched in *Endozoicomonas* (ASV_22), BD1-7 clade (ASV_2), and *Candidatus Hepatoplasma* (ASV_6), while *Candidatus Amoebophilus* (ASV_84), Proteobacteria_IS (ASV_7), BD7-8_IS (ASV_191), *Blattabacteriaceae*_IS (ASV_71) and *Endozoicomonas* (ASV_5) were differentially not enriched compared to the other three corals. In the case of *D. pertusum,* the SUP05 cluster (ASV_14) was differentially enriched, while we found that *Candidatus Hepatoplasma* (ASV_3 and 6) and the BD1-7 clade (ASV_2) were significantly less abundant when compared with the other target species. Moreover, we found similar patterns in *M. oculata,* where BD7-8_IS (ASV_77) was differentially enriched, and the BD1-7 clade (ASV_2) was underrepresented, contrary to what occurred in *A. hirsuta*. For *L. glaberrima,* we did not find any differentially abundant taxa **(Figure 5A)**.

In the microeukaryotic microbial community, we found differentially enriched Ichthyodinida-Clade-2_IS (ASVs 245, 502) in *A. hirsuta*, while we found differentially not enriched *Synophrya* (ASVs 82 and 122) and Ichthyodinida-Clade-2_IS (ASV 112). Comparing this with *M. oculata*, we found similarities, such as the differential presence of Ichthyodinida-Clade-2_IS (ASV 699). However, Ichthyodinida-Clade-2_IS (ASV_502) was differentially not enriched. For *D. pertusum*, we found Ichthyodinida-Clade-1_IS (ASV 520) and Ichthyodinida-Clade-2_IS (ASVs 671, 996, 357, and 438) were differentially not enriched. Finally, in *L. glaberrima* we found differentially enriched *Synophrya* (ASVs 82 and 122), in contrast to *A. hirsuta*, and we found differentially not enriched Ichthyodinida-Clade-2_IS (ASV 112) in contrast to *D. pertusum* **(Figure 5B)**.

## DISCUSSION

This study represents the first characterization of deep-sea coral microbiomes at Cabliers Reef, providing new insights into their structure and associations. Furthermore, based on the literature described up to this point, it explores the potential role of these microorganisms in the coral holobiont and its adaptation to the extreme conditions of Cabliers. Alpha diversity analysis revealed trends suggesting distinct ecological strategies among species. *A. hirsuta* presented a unique diversity profile, displaying consistently low prokaryotic diversity, while it showed the highest microeukaryotic diversity. These results are related to the dominance of a few prokaryotic taxa who are holding the main functional niches, and a varied microeukaryotic community that suggests its role in the degradation of organic matter captured by this soft coral species (Sánchez *et al*., 2019). *L. glaberrima* exhibited the most diverse prokaryotic community, suggesting a mature and stable microbiome that correlates with its longevity and is vital for maintaining physiological homeostasis throughout its lifetime (Roark *et al*., 2009). *D. pertusum* showed higher prokaryotic community diversity than *L. glaberrima* and *A. hirsuta*, but lower microeukaryotic community diversity. In addition, *D. pertusum* eukaryome diversity was similar to that shown by *M. oculata*. This last species, unlike the others, showed extremely high inter-individual variability, suggesting less predictable or more environmentally driven microbiomes than the other three corals. Moreover, this could suggest greater plasticity (Galand *et al*., 2020), enabling the coral to grow in different habitats and conditions, which may explain why it constitutes the majority of the coral population in Cabliers (Corbera *et al*., 2019).

Beta diversity analysis also confirmed that species identity was a determining factor in differences among microbial communities. We wanted to highlight the similarity in microbial community structure between *M. oculata* and *D. pertusum*, both scleractinian corals, suggesting that their aragonite-skeleton requirements and metabolic demands may also exert selective pressure on their associated microbes (Ianniello *et al*., 2025). Unlike the other species, *A. hirsuta* (Octocoral) is an outlier, suggesting a link between coral species and differences in their microbiomes.

Compositional analysis also provided insights into which key microorganisms shape these community structures and the roles they may play in each of our target species. While *Endozoicomonas* dominated the prokaryotic community in *M. oculata* (accounting for up for more than 40% of relative abundance), it was rarely found in association with the other corals. This genus is known for forming microbial aggregates associated with coral cells (CAMAs) and plays a vital role in group B vitamin metabolism and the sulfur cycle (Gotze *et al*., 2025a). These bacteria can metabolize sulfur compounds such as dimethylsulfoniopropionate (DMSP), converting it into dimethyl sulfide (DMS). In spite of the fact that DMSP is commonly associated with tropical corals, deep corals can also present this compound acting as an osmolyte and antioxidant in coral tissue (Kellogg and Pratte, 2021). In fact, a 16S rRNA meta-analysis of seven deep-sea coral species from the north-western Atlantic (Baltimore and Norfolk canyons) showed that *Endozoicomonas* was extremely rare (Kellogg, 2019). However, that meta-analysis did not include *M. oculata* or any Mediterranean coral. In a Mediterranean submarine canyon, *M. oculata* harbored temporally and spatially stable bacterial communities dominated by *Endozoicomonas* (Meistertzheim *et al*., 2016), which remained active after several months in aquaria (Galand *et al*., 2018). *Endozoicomonas*-dominated microbiomes have also been reported in deep-sea *Acanthogorgia* spp. and *D. pertusum* from some Atlantic sites (Kellogg and Pratte, 2021; Modolon *et al*., 2025). Thus, the dominance of *Endozoicomonas* in *M. oculata* at Cabliers seems to be a host-specific trait, at least in the Mediterranean, rather than an exception among deep-sea corals. Sulfur metabolism not only provides energy but also plays a defensive role through antioxidative mechanisms; thus, this prevalence suggests it may protect *M. oculata* from cellular stress under high hydrostatic pressure, thereby promoting active coral growth (Gotze *et al*., 2025b).

Another remarkable discovery is the high abundance of the cluster ASV_14 in *D. pertusum,* which accounts for up to 32.2% of its total prokaryotic community. These chemoautotrophic bacteria are common sulfur oxidants living in cold seeps. Their association with deep-sea corals suggest that the holobiont might have developed a facultative strategy to fix carbon independently of photosynthesis (Zhou *et al*., 2019; Vohsen *et al*., 2025). The coexistence with sulfur-oxidizing symbionts relationship could also explain the vigorous growth of these corals in Cabliers, by providing another source of organic matter beyond their heterotrophic diet based on zooplankton (Dede *et al*., 2022). Actually, a recent survey screening deep sea coral near cold seeps for Thioglobaceae (the family containing the SUP05 cluster) found ASVs associated with coral hosts at high relative abundance, ranging from 10 to 91%. These ASVs were specific to coral tissue and largely absent from surrounding water and sediment. Moreover, through metagenomics and transcriptomics they confirmed that the sulfur-oxidation and carbon-fixation pathways encoded by these symbionts were transcriptionally active (Vohsen *et al*., 2025).

Regarding *A. hirsuta*, our compositional analysis revealed it to be the least similar coral among the species in this study. It harbors significantly distinct taxa compared to the other corals, such as Mollicutes or BD1-7. Entomoplasmatales is an order within the class Mollicutes, so the genera *Candidatus* Hepatoplasma and *Candidatus* Oceanoplasma also belong to this group. Together, BD1-7 clade and Mollicutes account for more than 75.0% of the whole microbial community in the gorgonian species, suggesting an important role as core member of the holobiont and in the coral’s survival. These bacteria can degrade complex polysaccharides such as chitin or cellulose (van de Water *et al*., 2018). Therefore, BD1-7 clade bacteria likely decompose the crustaceans’ chitinous exoskeletons eaten by the corals, facilitating easier digestion and nutrient capture. This metabolic capability enables the coral to grow on volcanic mounds, where water currents constantly provide a steady supply of organic particles (Ho *et al*., 2025). Besides, Mollicutes’ significant presence is represented by the genera *Candidatus Hepatoplasma* and *Candidatus Oceanoplasma*; these bacteria cannot metabolize carbohydrates and instead rely on host-derived arginine degradation (Faburay and McVey, 2022). Thus, they could be providing bioavailable nitrogen to the coral. Moreover, the presence of *Hepatoplasma* is positively correlated with the abundance of the BD1-7 clade (van de Water *et al*., 2018), suggesting a complex, interactive microbiome network that optimizes nitrogen assimilation for the holobiont. Compared with current knowledge, our findings across cold-water gorgonians further support species- and depth-specific microbiomes: temperate *P. clavata* is dominated by Endozoicomonas, whereas the deep-sea congener *P. placomus* lacks it entirely, and *Primnoa pacifica/P. resedaeformis* show a similar shift from *Chlamydiae* or *Endozoicomonas* dominated shallow communities to far more diverse deep-sea ones (Goldsmith *et al*., 2018).

For the prokaryotic community *of L. glaberrima*, bubble plots and heat maps showed dominance of Proteobacteria, particularly the genus *Alteromonas* ASV_51 (6.9% read abundance). Corals secrete mucus that is enriched in polysaccharides, proteins, and lipids. This bacterial genus is associated with degrading organic matter and providing the coral with chemical defense by producing antimicrobial peptides or iron chelators (siderophores) (Mohamed *et al*., 2023). In addition, Bacteroidota *Candidatus Amoebophilus* was found (8.0% read abundance), could be used as an indicator of microeukaryotic activity and highlighting a multitrophic symbiotic cascade. This is because they seem to be obligate intracellular bacteria within protists (Schmitz-Esser *et al*., 2010). Amoebophilaceae sequences are commonly recovered from coral tissue, and these bacteria have been proposed to reside inside protists associated with the coral rather than within coral cells themselves (Apprill *et al*., 2016; Huggett and Apprill, 2019).

If we consider Cabliers’ microeukaryotes, they were less diverse than the prokaryotic microbiome. As expected, given the great depth conditions at Cabliers, we did not find *Symbiodiniaceae* or Corallicolida as main symbionts, as it is common in tropical corals. Instead, Ichthyodinida and secondarily Syndiniales (Amoebophryida) dominated the eukaryotic community. These are obligate parasites, and their presence suggests they may control microbiome balance, preventing other opportunistic taxa from proliferating. Recent studies suggest this prevalence could indicate an important role in carbon flux, forming a carbon-recycling engine that liberates organic matter through host cell lysis (Clarke *et al*., 2019; Anderson *et al*., 2024), providing an energy source for heterotrophic bacteria. Furthermore, Ichthyodinium was detected in all coral core microbiomes analyzed *(M. oculata*, *D. pertusum*, and *L. glaberrima*) but not in *A. hirsuta*. Metabarcoding detected it in both hard and soft corals, with high relative abundance in octocorals and black corals (del Campo *et al*., 2026). The consistent presence of this group of microorganisms in our studied corals and throughout the literature suggests it is not a coincidence. These microbes may represent a stable associate of the holobiont rather than an incidental colonizer, as they could be coral parasites or parasites of other protists (Bonacolta *et al*., 2024). Notably, Ichthyodinida were recently found significantly enriched in thermally resistant colonies of the Mediterranean gorgonian *Paramuricea clavata* (Bonacolta *et al*., 2024), suggesting that the relationship between this clade known for being parasitic, and the coral holobiont may not be strictly parasitic. This way, Ichthyodinida could contribute to processes as protection and stress modulation.

Concerning *L. glaberrima,* eukaryome analysis revealed a particularly distinct signature, with a clear dominance of the genera *Synophrya* (ASVs 82 and 122) and *Vampyrophrya* ASV_251, representing 41.5% and 8.2% of total read abundance, respectively, an association not observed in any other coral species in this study. These are apostome ciliates known to be associated with crustaceans; however, these results suggest that the mucus layer of *L. glaberrima,* or the bacteria within it, is being consumed by these microorganisms, thereby cleaning the coral’s superficial layer and enabling other protists to colonize the niche (Ohtsuka *et al*., 2015).

Once we documented specific taxa for each species, we next examined which microorganisms were involved and which were shared or not shared between species. Upset plots showed that, in both the prokaryotic and microeukaryotic communities, some ASVs were exclusive to each coral species, while others were shared. In addition to the taxa specific to each species, the four corals shared a set of microorganisms that may also help explain how they thrive in this environment. This core may be characteristic of the Cabliers region and help the reef thrive. This vitality is the result of the nutrient balance and recycling within the holobiont. The presence of *Nitrosopumilus* in all species suggests that nitrification is important for deep corals, as these microorganisms use the ammonium produced by their hosts as an energy source while maintaining detoxified coral tissues (Webb *et al*., 2025). In addition, *Endozoicomonas* has recently been shown to accumulate phosphorus (Gotze *et al*., 2025b); thus, because phosphorus is limited in the Mediterranean Sea (Pujo-Pay *et al*., 2011), *Endozoicomonas* could be essential for skeleton growth and ATP synthesis. Likewise, bacterial orders such as *Campylobacterales* ASV_54 (6.2% read abundance in *D. pertusum*) and *Spirochaetales* are significantly present in *A. hirsuta,* suggesting alternative energy sources, as they can oxidize reduced sulfur compounds. However, core microbiome analysis also showed exclusive ASVs (those detected in upset plots). ANCOM analysis confirmed that the BD1-7 clade (ASV_2) was differentially enriched only for *A. hirsuta* and differentially depleted for *M. oculata* and *D. pertusum*. For this last species, we also confirmed that the SUP05 cluster (ASV_14) was differentially enriched. We found similar patterns between *M. oculata* and *D. pertusum,* as they present absences of taxa that were present, or the other way around, in *A. hirsuta* (BD1-7 clade (ASV_2), *Candidatus Hepatoplasma* (ASV_3 and 6) or BD7-8_IS (ASV_77)). These results suggest that a core microbiome shared by the deep corals at Cabliers coexists with ASVs exclusive to each species. These species-specific taxa define a distinctive signature microbiome for each coral.

To sum up, the microbial diversity shown in this study reveals the richness of the cold-water coral reef in the Cabliers environment. The discovery provides insights of microbial metabolic niches suggests that losing a single coral species could reduce not only biodiversity complexity but also vital microbiome functions across the entire ecosystem, highlighting the importance of maintaining and protecting Cabliers cold-water coral reefs from anthropogenic activities.

## CONCLUSIONS

Our results show that host identity is the primary factor regulating the structure and composition of both prokaryotic and microeukaryotic communities across the four coral species studied at Cabliers, implying that losing any single coral species from the reef could entail a disproportionate loss of unique microbial functional niches. This host-driven structuring is further reflected in the divergence between the octocoral *Acanthogorgia hirsuta*, dominated by the chitin-degrading BD1-7 clade and by Mollicutes, the scleractinians *Madrepora oculata* and *Desmophyllum pertusum*, characterized by *Endozoicomonas* and the chemoautotrophic SUP05 cluster, respectively, and the black coral *Leiopathes glaberrima*, which harbored the most diverse prokaryotic community and a microeukaryotic community dominated by apostome ciliates, collectively suggesting that the chemical and physical nature of the coral skeleton constitutes a fundamental axis of microbial metabolic niche differentiation.

Several of the taxa identified here probably perform key functions for their hosts, such as sulfur oxidation and carbon fixation by the SUP05 cluster in *D. pertusum*, nutrient and sulfur cycling by *Endozoicomonas* in *M. oculata*, chitin degradation and nitrogen supply by the BD1-7 clade and Mollicutes in *A. hirsuta*, and nitrification by *Nitrosopumilus* in all four species. Because these microorganisms are interconnected through the exchange of metabolites, the loss of any of them could unbalance the holobiont and compromise the growth or survival of its coral host. Such losses are plausible under ocean warming, which in cold water corals (CWCs) alters the microbiome before host physiology declines and has been linked to pathogen-associated mortality (Chapron *et al*., 2021; Chemel *et al*., 2024). Since the four species studied here jointly build the three-dimensional framework of Cabliers, the decline of a single one would alter the habitat and the microbial functions it supports, jeopardizing the stability of the whole reef. Characterizing these interactions and their functions, for instance through metagenomics and metatranscriptomics, is therefore essential to protect the only actively growing CWC reef in the Mediterranean.

## Supporting information

Data S1. 16S rRNA ASV count table with taxonomy

Data S2. 18S rRNA ASV count table with taxonomy

Data S3. Sample metadata

## ACKNOWLEDGMENTS

The authors acknowledge the Spanish National PROTEUS Project (MCIN/AEI PID2021-122836OB-I00) and thank the crew and scientific team of the PROTEUS cruise expedition for sample collection at Cabliers Coral Mound. Sequencing was performed at the Integrated Microbiome Resource (IMR), Dalhousie University, Canada. This work was supported by JAE Intro ICU 2025 CSIC’s program. The authors declare that no third-party figures or copyrighted material requiring permission have been reproduced in this manuscript.

## AUTHOR CONTRIBUTIONS

Aimar Ardanaz-Ullate: Conceptualization, Methodology, Formal analysis, Investigation, Visualization, Writing original draft. Agazzi M: Methodology, DNA/RNA extractions. Lo Iacono C: Conceptualization, Sampling, Funding acquisition. del Campo J: Conceptualization, Supervision, Writing review & editing, Funding acquisition.

## DATA AVAILABILITY STATEMENT

The 16S and 18S rRNA gene sequence data generated in this study have been deposited in the NCBI Sequence Read Archive (SRA) under BioProject accession number **PRJNA1480519**. The R pipeline is provided in https://github.com/delCampoLab/ardanaz_cabliers_pipeline. All other data are available from the corresponding author upon reasonable request.

## FUNDER INFORMATION DECLARED

The project that gave rise to these results received the support of a fellowship from Consejo Superior de Investigaciones Científicas, https://ror.org/02gfc7t72, JAE Intro ICU 2025, JAEICU_25_00113 to AAU.

This project was supported by Grant PID2023-152522NB-I00, funded by MICIU/AEI/10.13039/501100011033 and by ERDF, EU provided to JdC.

## REFERENCES

Andersen, K.S., Kirkegaard, R.H., Karst, S.M., and Albertsen, M. (2018) ampvis2: an R package to analyse and visualise 16S rRNA amplicon data. bioRxiv 299537. 10.1101/299537

Anderson, S.R., Blanco-Bercial, L., Carlson, C.A., and Harvey, E.L. (2024) Role of Syndiniales parasites in depth-specific networks and carbon flux in the oligotrophic ocean. ISME Commun 4: ycae014.

Apprill, A., McNally, S., Parsons, R., and Weber, L. (2015) Minor revision to V4 region SSU rRNA 806R gene primer greatly increases detection of SAR11 bacterioplankton. Aquat Microb Ecol 75: 129–137.

Apprill, A., Weber, L.G., and Santoro, A.E. (2016) Distinguishing between microbial habitats unravels ecological complexity in coral microbiomes. mSystems 1: e00143–16.

Bo, M., Cerrano, C., Canese, S., Salvati, E., Angiolillo, M., Santangelo, G., and Bavestrello, G. (2014) The coral assemblages of an off-shore deep Mediterranean rocky bank (NW Sicily, Italy). Mar Ecol 35: 332–342.

Bonacolta, A.M., Miravall, J., Gómez-Gras, D., Ledoux, J.-B., López-Sendino, P., Garrabou, J., et al. (2024) Differential apicomplexan presence predicts thermal stress mortality in the Mediterranean coral *Paramuricea clavata*. Environ Microbiol 26: e16548.

Callahan, B.J., McMurdie, P.J., Rosen, M.J., Han, A.W., Johnson, A.J.A., and Holmes, S.P. (2016a) DADA2: high-resolution sample inference from Illumina amplicon data. Nat Methods 13: 581–583.

Callahan, B.J., Sankaran, K., Fukuyama, J.A., McMurdie, P.J., and Holmes, S.P. (2016b) Bioconductor workflow for microbiome data analysis: from raw reads to community analyses. F1000Research 5: 1492.

Chapron, L., Galand, P.E., Pruski, A.M., Peru, E., Vétion, G., Robin, S., and Lartaud, F. (2021) Resilience of cold-water coral holobionts to thermal stress. Proc R Soc B 288: 20212117.

Chemel, M., Peru, E., Binsarhan, M., Logares, R., Lartaud, F., and Galand, P.E. (2024) Cold-water coral mortality under ocean warming is associated with pathogenic bacteria. Environ Microbiome 19: 76.

Clarke, L.J., Bestley, S., Bissett, A., and Deagle, B.E. (2019) A globally distributed Syndiniales parasite dominates the Southern Ocean micro-eukaryote community near the sea-ice edge. ISME J 13: 734–737.

Comeau, A.M., Li, W.K.W., Tremblay, J.-É., Carmack, E.C., and Lovejoy, C. (2011) Arctic Ocean microbial community structure before and after the 2007 record sea ice minimum. PLoS One 6: e27492.

Corbera, G., Lo Iacono, C., Gràcia, E., Grinyó, J., Pierdomenico, M., Huvenne, V.A.I., et al. (2019) Ecological characterisation of a Mediterranean cold-water coral reef: Cabliers Coral Mound Province (Alboran Sea, western Mediterranean). Prog Oceanogr 175: 245–262.

Corbera, G., Lo Iacono, C., Simarro, G., Grinyó, J., Ambroso, S., Huvenne, V.A.I., et al. (2022) Local-scale feedbacks influencing cold-water coral growth and subsequent reef formation. Sci Rep 12: 20389.

Dede, B., Hansen, C.T., Neuholz, R., Schnetger, B., Kleint, C., Walker, S., et al. (2022) Niche differentiation of sulfur-oxidizing bacteria (SUP05) in submarine hydrothermal plumes. ISME J 16: 1479–1490.

del Campo, J., Pons, M.J., Herranz, M., Wakeman, K.C., del Valle, J., Vermeij, M.J.A., et al. (2019) Validation of a universal set of primers to study animal-associated microeukaryotic communities. Environ Microbiol 21: 3855–3861.

del Campo, J., Bonacolta, A.M., Weiler, B.A., Knowles, B., Apprill, A., Fox, M.D., et al. (2026) Global diversity and distribution of coral-associated protists. bioRxiv 2026.01.22.700988. 10.64898/2026.01.22.700988

Faburay, B., and McVey, D.S. (2022) Mollicutes. In Veterinary Microbiology. Hoboken, NJ: John Wiley & Sons, pp. 364–376.

Galand, P.E., Chapron, L., Meistertzheim, A.-L., Peru, E., and Lartaud, F. (2018) The effect of captivity on the dynamics of active bacterial communities differs between two deep-sea coral species. Front Microbiol 9: 2565.

Galand, P.E., Remize, M., Meistertzheim, A.-L., Pruski, A.M., Peru, E., Suhrhoff, T.J., et al. (2020) Diet shapes cold-water corals bacterial communities. Environ Microbiol 22: 354–368.

Gloor, G.B., Macklaim, J.M., Pawlowsky-Glahn, V., and Egozcue, J.J. (2017) Microbiome datasets are compositional: and this is not optional. Front Microbiol 8: 2224.

Goldsmith, D.B., Kellogg, C.A., Morrison, C.L., Gray, M.A., Stone, R.P., Waller, R.G., et al. (2018) Comparison of microbiomes of cold-water corals *Primnoa pacifica* and *Primnoa resedaeformis*, with possible link between microbiome composition and host genotype. Sci Rep 8: 12383.

Gori, A., Wienberg, C., Grinyó, J., Taviani, M., Hebbeln, D., Lo Iacono, C., et al. (2023) Life and death of cold-water corals across the Mediterranean Sea. In Cold-Water Coral Reefs of the World. Cordes, E., and Mienis, F. (eds). Cham: Springer International Publishing, pp. 171–197.

Gotze, C.R., Dungan, A.M., van de Meene, A.M.L., Damjanovic, K., Philip, G.K., Maire, J., et al. (2025a) Differential aggregation patterns of *Endozoicomonas* within tissues of the coral *Acropora loripes*. ISME J 19: wraf059.

Gotze, C.R., Tandon, K., Philip, G.K., Dungan, A.M., Maire, J., Høj, L., et al. (2025b) Genomic prediction of symbiotic interactions between two *Endozoicomonas* clades and their coral host, *Acropora loripes*. bioRxiv 2025.02.16.638547. 10.1101/2025.02.16.638547

Guillou, L., Bachar, D., Audic, S., Bass, D., Berney, C., Bittner, L., et al. (2013) The Protist Ribosomal Reference database (PR2): a catalog of unicellular eukaryote small sub-unit rRNA sequences with curated taxonomy. Nucleic Acids Res 41: D597–D604.

Henry, L.-A., and Roberts, J.M. (2016) Global biodiversity in cold-water coral reef ecosystems. In Marine Animal Forests: The Ecology of Benthic Biodiversity Hotspots. Rossi, S., Bramanti, L., Gori, A., and Orejas, C. (eds). Cham: Springer International Publishing, pp. 1–21.

Hernandez-Agreda, A., Gates, R.D., and Ainsworth, T.D. (2017) Defining the core microbiome in corals’ microbial soup. Trends Microbiol 25: 125–140.

Ho, J.Y., Koh, X.Q., Kang, D.Y., Low, A., Hu, D., Haryono, M.A.S., et al. (2025) Discovery of a phylogenetically novel tropical marine Gammaproteobacteria elucidated from assembled genomes and the proposed transfer of the genus *Umboniibacter* from the family Cellvibrionaceae to Umboniibacteraceae fam. nov. Front Microbiol 16: 1437936.

Huggett, M.J., and Apprill, A. (2019) Coral microbiome database: integration of sequences reveals high diversity and relatedness of coral-associated microbes. Environ Microbiol Rep 11: 372–385.

Ianniello, C.F., McAlister, J.S., Ferrier-Pagès, C., and Rotjan, R. (2025) A review of the current knowledge of the flow of carbon and energy in scleractinian corals. Coral Reefs 44: 2191–2233.

Kellogg, C.A. (2019) Microbiomes of stony and soft deep-sea corals share rare core bacteria. Microbiome 7: 90.

Kellogg, C.A., and Pratte, Z.A. (2021) Unexpected diversity of *Endozoicomonas* in deep-sea corals. Mar Ecol Prog Ser 673: 1–15.

Lauria, V., Massi, D., Fiorentino, F., Milisenda, G., and Cillari, T. (2021) Habitat suitability mapping of the black coral *Leiopathes glaberrima* to support conservation of vulnerable marine ecosystems. Sci Rep 11: 15661.

Lin, H., and Peddada, S.D. (2020) Analysis of compositions of microbiomes with bias correction. Nat Commun 11: 3514.

Lin, H., and Peddada, S.D. (2024) Multigroup analysis of compositions of microbiomes with covariate adjustments and repeated measures. Nat Methods 21: 83–91.

Lo Iacono, C., Gràcia, E., Ranero, C.R., Emelianov, M., Huvenne, V.A.I., Bartolomé, R., et al. (2014) The West Melilla cold water coral mounds, Eastern Alboran Sea: morphological characterization and environmental context. Deep Sea Res Part II Top Stud Oceanogr 99: 316–326.

Lo Iacono, C., Savini, A., Huvenne, V.A.I., and Gràcia, E. (2019) Habitat mapping of cold-water corals in the Mediterranean Sea. In Mediterranean Cold-Water Corals: Past, Present and Future. Orejas, C., and Jiménez, C. (eds). Cham: Springer International Publishing, pp. 157–171.

Martin, M. (2011) Cutadapt removes adapter sequences from high-throughput sequencing reads. EMBnet J 17: 10–12.

Massi, D., Vitale, S., Titone, A., Milisenda, G., Gristina, M., and Fiorentino, F. (2018) Spatial distribution of the black coral Leiopathes glaberrima (Esper, 1788) (Antipatharia: Leiopathidae) in the Mediterranean: a prerequisite for protection of Vulnerable Marine Ecosystems (VMEs). Eur Zool J 85: 169–178.

Mayer, K.-I., Colin, L., Voolstra, C.R., and Cárdenas, A. (2025) Ecosystem microbiology of coral reefs. In Coral Reef Microbiome. Peixoto, R.S., and Voolstra, C.R. (eds). Coral Reefs of the World, Vol. 20. Cham: Springer International Publishing, pp. 131–144.

McMurdie, P.J., and Holmes, S. (2013) phyloseq: an R package for reproducible interactive analysis and graphics of microbiome census data. PLoS One 8: e61217.

Meistertzheim, A.-L., Lartaud, F., Arnaud-Haond, S., Kalenitchenko, D., Bessalam, M., Le Bris, N., and Galand, P.E. (2016) Patterns of bacteria-host associations suggest different ecological strategies between two reef building cold-water coral species. Deep Sea Res Part I Oceanogr Res Pap 114: 12–22.

Modolon, F., Garritano, A.N., Hill, L.J., Duarte, G., Bendia, A., de Moura, R., et al. (2025) Putative promiscuous symbionts in deep-sea corals and crinoids may contribute to nitrogen cycling. Microbiome 13: 234.

Mohamed, A.R., Ochsenkühn, M.A., Kazlak, A.M., Moustafa, A., and Amin, S.A. (2023) The coral microbiome: towards an understanding of the molecular mechanisms of coral-microbiota interactions. FEMS Microbiol Rev 47: fuad005.

Ohtsuka, S., Suzaki, T., Kanazawa, A., and Ando, M. (2015) Biology of symbiotic apostome ciliates: their diversity and importance in the aquatic ecosystems. In Marine Protists: Diversity and Dynamics. Ohtsuka, S., Suzaki, T., Horiguchi, T., Suzuki, N., and Not, F. (eds). Tokyo: Springer Japan, pp. 441–463.

Oksanen, J., Simpson, G.L., Blanchet, F.G., Kindt, R., Legendre, P., Minchin, P.R., et al. (2025) vegan: community ecology package. R package. URL https://CRAN.R-project.org/package=vegan

Orejas, C., and Jiménez, C. (eds) (2019) Mediterranean Cold-Water Corals: Past, Present and Future. Understanding the Deep-Sea Realms of Coral. Coral Reefs of the World, Vol. 9. Cham: Springer International Publishing.

Parada, A.E., Needham, D.M., and Fuhrman, J.A. (2016) Every base matters: assessing small subunit rRNA primers for marine microbiomes with mock communities, time series and global field samples. Environ Microbiol 18: 1403–1414.

Pujo-Pay, M., Conan, P., Oriol, L., Cornet-Barthaux, V., Falco, C., Ghiglione, J.-F., et al. (2011) Integrated survey of elemental stoichiometry (C, N, P) from the western to eastern Mediterranean Sea. Biogeosciences 8: 883–899.

Quast, C., Pruesse, E., Yilmaz, P., Gerken, J., Schweer, T., Yarza, P., et al. (2013) The SILVA ribosomal RNA gene database project: improved data processing and web-based tools. Nucleic Acids Res 41: D590–D596.

Raimundo, J., Vale, C., Caetano, M., Anes, B., Carreiro-Silva, M., Martins, I., et al. (2013) Element concentrations in cold-water gorgonians and black coral from Azores region. Deep Sea Res Part II Top Stud Oceanogr 98: 129–136.

Roark, E.B., Guilderson, T.P., Dunbar, R.B., Fallon, S.J., and Mucciarone, D.A. (2009) Extreme longevity in proteinaceous deep-sea corals. Proc Natl Acad Sci USA 106: 5204–5208.

Roberts, J.M., and Cairns, S.D. (2014) Cold-water corals in a changing ocean. Curr Opin Environ Sustain 7: 118–126.

Roberts, J.M., Wheeler, A.J., and Freiwald, A. (2006) Reefs of the deep: the biology and geology of cold-water coral ecosystems. Science 312: 543–547.

Roberts, J.M., Wheeler, A.J., Freiwald, A., and Cairns, S.D. (2009) Cold-Water Corals: The Biology and Geology of Deep-Sea Coral Habitats. Cambridge: Cambridge University Press.

Sánchez, J.A., Dueñas, L.F., Rowley, S.J., Gonzalez-Zapata, F.L., Vergara, D.C., Montaño-Salazar, S.M., et al. (2019) Gorgonian corals. In Mesophotic Coral Ecosystems. Loya, Y., Puglise, K.A., and Bridge, T.C.L. (eds). Cham: Springer International Publishing, pp. 729–747.

Sartoretto, S., and Zibrowius, H. (2018) Note on new records of living Scleractinia and Gorgonaria between 1700 and 2200 m depth in the western Mediterranean Sea. Mar Biodivers 48: 689–694.

Schmitz-Esser, S., Tischler, P., Arnold, R., Montanaro, J., Wagner, M., Rattei, T., and Horn, M. (2010) The genome of the amoeba symbiont ‘*Candidatus* Amoebophilus asiaticus’ reveals common mechanisms for host cell interaction among amoeba-associated bacteria. J Bacteriol 192: 1045–1057.

United Nations (2017) Cold-water corals. In The First Global Integrated Marine Assessment: World Ocean Assessment I. Cambridge: Cambridge University Press, pp. 803–816.

van de Water, J.A.J.M., Allemand, D., and Ferrier-Pagès, C. (2018) Host-microbe interactions in octocoral holobionts: recent advances and perspectives. Microbiome 6: 64.

van Oppen, M.J.H., and Blackall, L.L. (2019) Coral microbiome dynamics, functions and design in a changing world. Nat Rev Microbiol 17: 557–567.

Vohsen, S.A., Gruber-Vodicka, H.R., Osman, E.O., Saxton, M.A., Joye, S.B., Dubilier, N., et al. (2025) Deep-sea corals near cold seeps associate with sulfur-oxidizing chemoautotrophs in the family *Ca.* Thioglobaceae. Microbiome 13: 232.

Voolstra, C.R., Raina, J.-B., Dörr, M., Cárdenas, A., Pogoreutz, C., Silveira, C.B., et al. (2024) The coral microbiome in sickness, in health and in a changing world. Nat Rev Microbiol 22: 460–475.

Webb, A.L., Oudova-Rivera, B., Ward, M., Carpenter, L.J., Lehtovirta-Morley, L.E., and Chance, R. (2025) The ammonia oxidising archaeon *Nitrosopumilus maritimus* does not alter iodine oxidation state in oxic seawater. Environ Microbiol Rep 17: e70168.

Weinnig, A.M., Aunins, A.W., Salamone, V., Quattrini, A.M., Nizinski, M.S., and Morrison, C.L. (2024) Population structure of *Desmophyllum pertusum* found along the United States eastern continental margin. BMC Res Notes 17: 326.

Wickham, H., Averick, M., Bryan, J., Chang, W., McGowan, L.D., François, R., et al. (2019) Welcome to the Tidyverse. J Open Source Softw 4: 1686.

Zhou, K., Zhang, R., Sun, J., Zhang, W., Tian, R.M., Chen, C., et al. (2019) Potential interactions between clade SUP05 sulfur-oxidizing bacteria and phages in hydrothermal vent sponges. Appl Environ Microbiol 85: e00992–19.

