## Supplementary Material (Tables S1-S3, Figures S1-S5) for "Host identity drives distinct prokaryotic and microeukaryotic microbiomes across four cold-water coral species in a near-pristine Mediterranean reef"

#### SUPPLEMENTARY TABLES

**Table S1.** Number of sequencing reads retained per sample at key processing steps for prokaryotic analysis. Input refers to raw reads, filtered to reads after quality filtering, denoisedF to forward noise-corrected reads, denoisedR to noise-corrected reverse reads, merged to merged reads, and nonchim to final reads retained after chimera removal.

| Sample | input | filtered | denoisedF | denoisedR | merged | nonchim |
|---|---|---|---|---|---|---|
| deep_1 | 58213 | 57983 | 57725 | 57759 | 57327 | 56313 |
| deep_100 | 4073 | 2526 | 2465 | 2492 | 2413 | 2409 |
| deep_101 | 55669 | 55161 | 54976 | 54957 | 54762 | 54745 |
| deep_102 | 47472 | 47109 | 46948 | 46950 | 46718 | 46706 |
| deep_103 | 56183 | 55740 | 55470 | 55510 | 55185 | 55158 |
| deep_11 | 32821 | 32589 | 31587 | 31802 | 30047 | 29505 |
| deep_12 | 88896 | 88189 | 84404 | 85478 | 80433 | 79374 |
| deep_2 | 42977 | 42706 | 42350 | 42381 | 41830 | 41330 |
| deep_26 | 57514 | 57161 | 56974 | 56996 | 56701 | 56080 |
| deep_3 | 57671 | 57389 | 56829 | 56845 | 56013 | 55360 |
| deep_30 | 71443 | 71071 | 70972 | 70983 | 70789 | 69861 |
| deep_31 | 22238 | 22099 | 22033 | 22039 | 21937 | 21749 |
| deep_33 | 2505 | 881 | 843 | 860 | 826 | 826 |
| deep_34 | 2409 | 2008 | 1982 | 1985 | 1957 | 1956 |
| deep_35 | 3102 | 2524 | 2498 | 2508 | 2476 | 2476 |
| deep_37 | 6905 | 6736 | 6719 | 6719 | 6384 | 6384 |
| deep_38 | 12982 | 12783 | 12766 | 12768 | 12751 | 12747 |
| deep_39 | 4844 | 4779 | 4756 | 4755 | 4731 | 4730 |
| deep_46 | 19421 | 16417 | 16357 | 16375 | 16297 | 16221 |
| deep_47 | 37881 | 36315 | 35613 | 35702 | 34573 | 34446 |
| deep_48 | 5943 | 1921 | 1859 | 1885 | 1817 | 1811 |
| deep_49 | 17514 | 17319 | 17231 | 17244 | 17038 | 17020 |
| deep_5 | 2925 | 2748 | 2701 | 2722 | 2667 | 2665 |
| deep_50 | 3992 | 3952 | 3916 | 3916 | 3894 | 3894 |
| deep_51 | 8615 | 8548 | 8508 | 8515 | 8469 | 8469 |
| deep_53 | 103172 | 102564 | 101805 | 101969 | 100854 | 97402 |
| deep_54 | 85867 | 85289 | 84935 | 85008 | 84389 | 82918 |
| deep_55 | 70532 | 70052 | 69695 | 69711 | 69110 | 67740 |
| deep_57 | 70990 | 70194 | 70065 | 70071 | 69889 | 69879 |
| deep_58 | 34266 | 34004 | 33959 | 33954 | 33912 | 33912 |

| | | | | | | |
|---|---|---|---|---|---|---|
| deep_59 | 27634 | 27373 | 27325 | 27305 | 27258 | 27254 |
| deep_6 | 2513 | 2419 | 2367 | 2363 | 2300 | 2298 |
| deep_7 | 5087 | 4349 | 4270 | 4271 | 4173 | 4170 |
| deep_71 | 49421 | 49041 | 48984 | 48993 | 48943 | 48943 |
| deep_72 | 34176 | 33968 | 33913 | 33918 | 33859 | 33859 |
| deep_73 | 36649 | 36360 | 36327 | 36312 | 36272 | 36272 |
| deep_9 | 40896 | 40522 | 39489 | 39689 | 38056 | 37879 |
| deep_98 | 34331 | 33341 | 32788 | 32760 | 31748 | 31714 |
| deep_99 | 36640 | 35710 | 34580 | 34666 | 32794 | 32756 |

**Table S2.** Number of sequencing reads retained per sample at key processing steps for microeukaryotic analysis. Input refers to raw reads, filtered to reads after quality filtering, denoisedF to forward noise corrected reads, denoisedR to noise corrected reverse reads, merged to merged reads, and nonchim to final reads retained after chimera removal.

| Sample | input | filtered | denoisedF | denoisedR | merged | nonchim |
|---|---|---|---|---|---|---|
| deep_1 | 160228 | 158375 | 158157 | 158293 | 154745 | 148564 |
| deep_100 | 133896 | 132996 | 132979 | 132985 | 131718 | 131413 |
| deep_101 | 130977 | 126927 | 126878 | 126886 | 125860 | 125640 |
| deep_102 | 150021 | 145362 | 145243 | 145318 | 143833 | 136704 |
| deep_103 | 134310 | 129948 | 129868 | 129881 | 128609 | 128072 |
| deep_11 | 107421 | 106291 | 105992 | 105924 | 101655 | 68366 |
| deep_12 | 95188 | 80060 | 79066 | 79567 | 74219 | 60104 |
| deep_2 | 142492 | 141576 | 141418 | 141494 | 138662 | 133713 |
| deep_26 | 138174 | 137191 | 136934 | 137134 | 134444 | 129971 |
| deep_3 | 164197 | 162713 | 162437 | 162592 | 158741 | 151896 |
| deep_30 | 126802 | 125945 | 125724 | 125888 | 123563 | 120285 |
| deep_31 | 151998 | 150924 | 150607 | 150818 | 147614 | 142855 |
| deep_33 | 113670 | 112905 | 112877 | 112886 | 111837 | 111509 |
| deep_34 | 111686 | 110311 | 110301 | 110307 | 109111 | 108956 |
| deep_35 | 128618 | 127562 | 127529 | 127549 | 126171 | 125153 |
| deep_37 | 128724 | 127928 | 127861 | 127906 | 126140 | 122148 |
| deep_38 | 94547 | 93494 | 93456 | 93487 | 92011 | 91619 |
| deep_39 | 117194 | 116644 | 116586 | 116628 | 115114 | 114194 |
| deep_46 | 151556 | 150610 | 150551 | 150583 | 149085 | 146887 |
| deep_47 | 122696 | 116954 | 116688 | 116794 | 113174 | 98177 |

| | | | | | | |
|---|---|---|---|---|---|---|
| deep_48 | 129872 | 129246 | 129222 | 129230 | 128147 | 127473 |
| deep_49 | 118852 | 117793 | 117737 | 117749 | 116139 | 115467 |
| deep_5 | 130946 | 129918 | 129876 | 129901 | 128609 | 128182 |
| deep_50 | 116506 | 115469 | 115448 | 115455 | 113864 | 113617 |
| deep_51 | 124291 | 123467 | 123395 | 123434 | 121853 | 121314 |
| deep_53 | 128211 | 126661 | 126393 | 126492 | 122544 | 115039 |
| deep_54 | 114837 | 113850 | 113681 | 113767 | 110026 | 103334 |
| deep_55 | 148174 | 146557 | 146262 | 146405 | 141597 | 130947 |
| deep_57 | 133926 | 132396 | 132326 | 132359 | 130585 | 128950 |
| deep_58 | 115681 | 114969 | 114888 | 114920 | 113562 | 111469 |
| deep_59 | 116218 | 115479 | 115435 | 115450 | 114308 | 113250 |
| deep_6 | 82509 | 81794 | 81776 | 81790 | 80920 | 80622 |
| deep_7 | 123971 | 122560 | 122545 | 122543 | 120946 | 120828 |
| deep_71 | 95452 | 91383 | 91336 | 91350 | 90419 | 88386 |
| deep_72 | 122681 | 116808 | 116754 | 116753 | 115211 | 114530 |
| deep_73 | 128758 | 122776 | 122729 | 122758 | 121148 | 120491 |
| deep_9 | 128743 | 124518 | 124169 | 124228 | 119845 | 90681 |
| deep_98 | 118559 | 117856 | 117769 | 117726 | 116188 | 115180 |
| deep_99 | 119262 | 118587 | 118451 | 118525 | 116752 | 115132 |

**Table S3.** Number of reads retained per sample after key processing steps of filtering the Phyloseq object (quality filtering, abundance filtering, unwanted sequences filtering, and decontamination).

| **16S rRNA gene** | | **18S rRNA gene** | |
|---|---|---|---|
| Sample_ID | Reads | Sample_ID | Reads |
| deep_1 | 55676 | deep_1 | 446 |
| deep_2 | 40150 | deep_2 | 341 |
| deep_3 | 53691 | deep_3 | 503 |
| deep_5 | 2526 | deep_5 | 302 |
| deep_6 | 2192 | deep_6 | 232 |
| deep_7 | 3942 | deep_7 | 103 |
| deep_9 | 34611 | deep_9 | 578 |

| | | | |
|---|---|---|---|
| deep_11 | 28712 | deep_11 | 174 |
| deep_12 | 74915 | deep_12 | 1153 |
| deep_26 | 55351 | deep_26 | 666 |
| deep_30 | 69526 | deep_30 | 260 |
| deep_31 | 21573 | deep_31 | 980 |
| deep_33 | 738 | deep_33 | 213 |
| deep_34 | 1913 | deep_34 | 120 |
| deep_35 | 2353 | deep_35 | 672 |
| deep_37 | 3210 | deep_37 | 229 |
| deep_38 | 10688 | deep_38 | 90 |
| deep_39 | 4521 | deep_39 | 348 |
| deep_46 | 15981 | deep_46 | 1362 |
| deep_47 | 32263 | deep_47 | 565 |
| deep_48 | 1692 | deep_48 | 382 |
| deep_49 | 16884 | deep_49 | 162 |
| deep_50 | 3152 | deep_50 | 79 |
| deep_51 | 8406 | deep_51 | 102 |
| deep_53 | 95901 | deep_53 | 685 |
| deep_54 | 82417 | deep_54 | 454 |
| deep_55 | 67105 | deep_55 | 616 |
| deep_57 | 2010 | deep_57 | 500 |
| deep_58 | 845 | deep_58 | 727 |
| deep_59 | 939 | deep_59 | 213 |
| deep_71 | 641 | deep_71 | 1284 |
| deep_72 | 952 | deep_72 | 197 |
| deep_73 | 365 | deep_73 | 210 |
| deep_98 | 29451 | deep_98 | 135 |
| deep_99 | 30445 | deep_99 | 212 |
| deep_100 | 2012 | deep_100 | 271 |
| deep_101 | 7478 | deep_101 | 66 |
| deep_102 | 7126 | deep_102 | 97 |
| deep_103 | 13127 | deep_103 | 172 |

### SUPPLEMENTARY FIGURES

**Figure S1.** Location of the Cabliers Coral Mound Province in the eastern Alboran Sea. The reef extends approximately 6 km along the northern flank of a volcanic ridge at depths of 250–710 m, between the southern coast of Spain and Morocco. Atlantic water inflow and complex ridge topography generate enhanced currents that sustain coral growth. Inset map shows the regional context within the western Mediterranean.

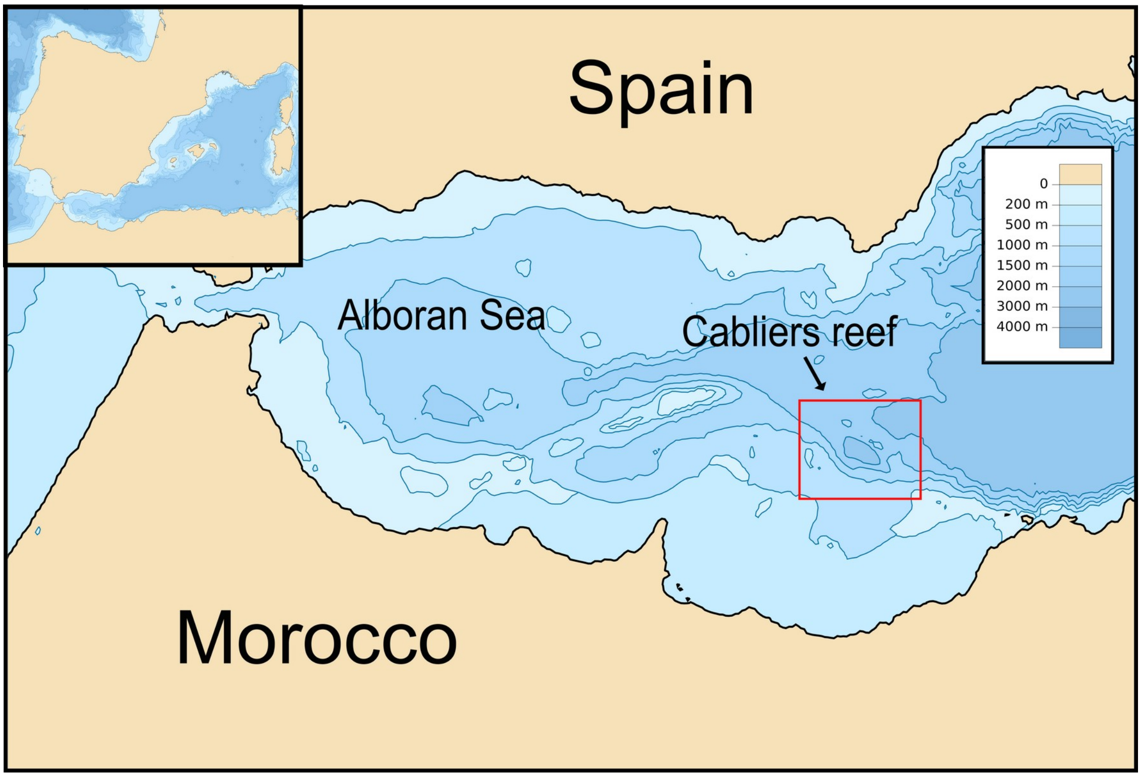


**Figure S2.** Bioinformatic pipeline overview. Raw Illumina MiSeq reads were quality-filtered and trimmed (Cutadapt), processed through the DADA2 pipeline for ASV inference and taxonomic assignment (SILVA v138 for 16S; PR2 v4.14.0 for 18S), and integrated into phyloseq for downstream ecological analyses. Primers: **A** 572F, **B** E1009R, **C** 515F, **D** 806RB.

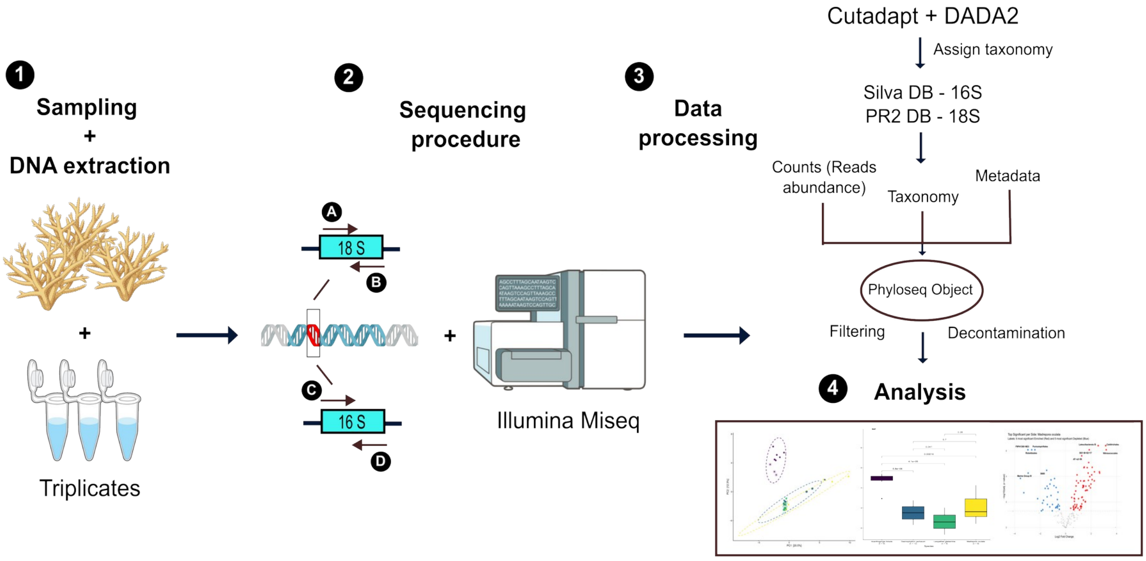

**Figure S3.** ANOSIM with 999 permutations was used to test for significant differences within the four different coral species for **A** prokaryotic and **B** microeukaryotic communities.

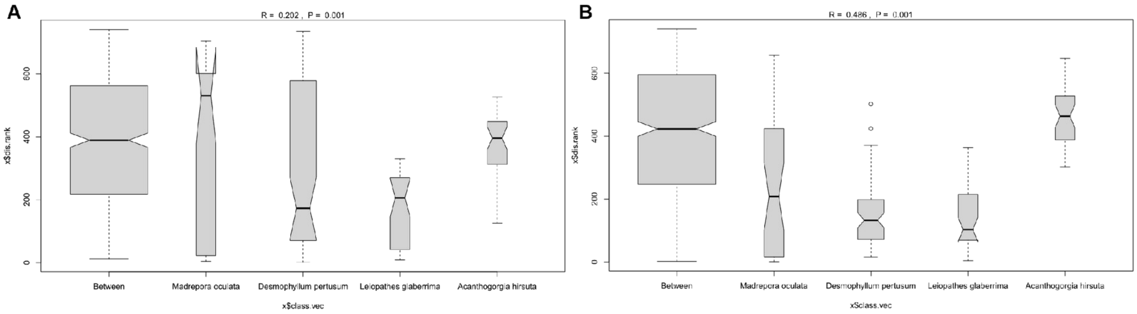


**Figure S4.** ANOSIM with 999 permutations was used to test for significant differences within the different dives for **A** prokaryotic and **B** microeukaryotic community.

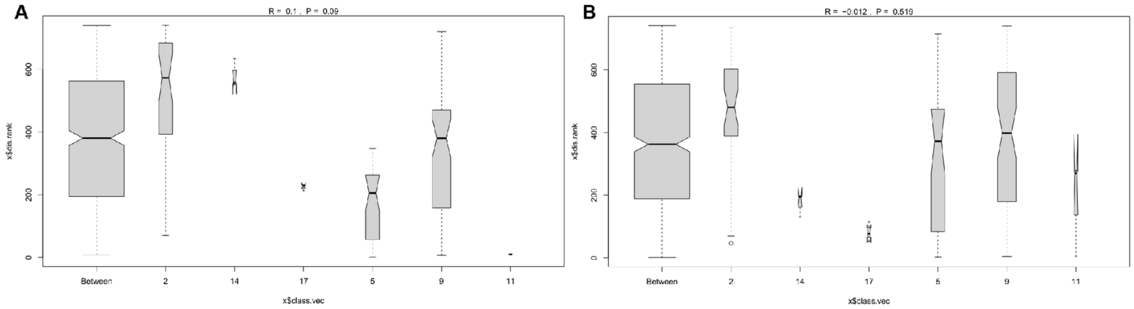


**Figure S5.** Alpha diversities from **A** prokaryotic and **B** microeukaryotic communities associated to the different dives. Showing significant Shannon alpha diversity community differences between some of the dives by pairwise comparisons, considering significant when (p≤0.05).

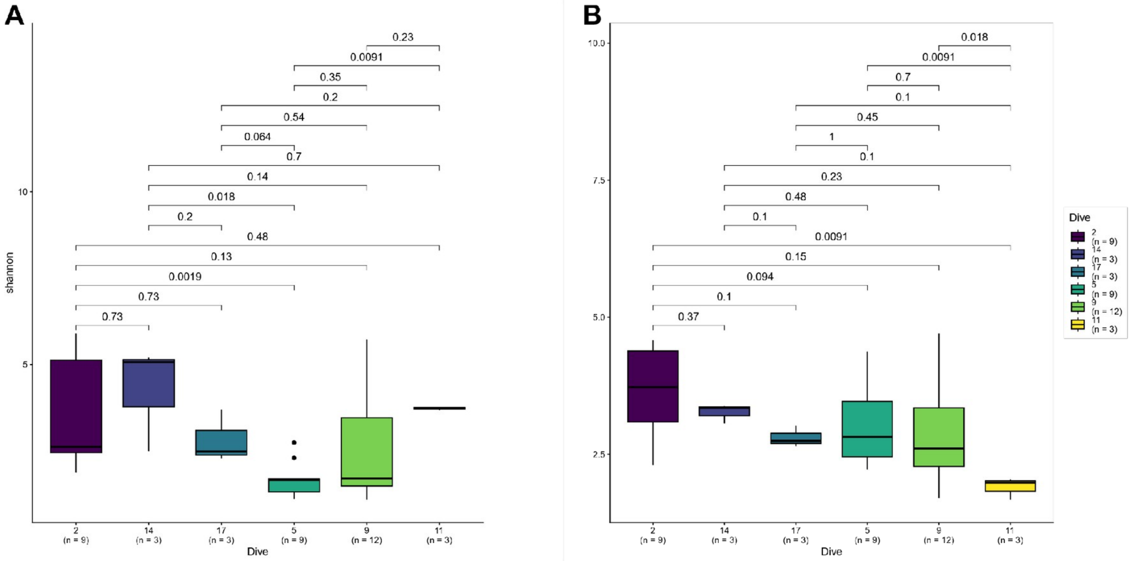
